# Investigating the Influence of Anti-Seizure Medications on Aperiodic EEG Activity

**DOI:** 10.1101/2025.11.02.686141

**Authors:** Marissa M. Holden, Isabella Premoli, Scott R. Clark, Mark P. Richardson, Mitchell R. Goldsworthy, Nigel C. Rogasch

## Abstract

**Objective:** Electroencephalography (EEG) signals comprise both oscillatory (periodic) and non-oscillatory (aperiodic) components. Aperiodic activity forms the 1/f-like background of the EEG power spectrum and can be characterised by parameters describing the offset and slope (1/f exponent). This study examined the effects of two antiseizure medications (ASMs), which reduce cortical excitability through different mechanisms of action, on aperiodic and periodic EEG activity.

**Methods:** Resting EEG was recorded with eyes open and closed from 13 healthy male volunteers at baseline and two hours after administration of lamotrigine (300 mg), levetiracetam (3000 mg), or placebo. Power spectra were computed using Welch’s method. Aperiodic parameters were estimated using the specparam algorithm, and periodic activity was quantified after subtraction of the aperiodic component.

**Results:** In the eyes-open condition, lamotrigine significantly reduced both aperiodic offset and exponent relative to placebo, consistent with a flattening of the aperiodic spectrum, whereas levetiracetam reduced the exponent without significantly altering the offset. Neither drug significantly altered aperiodic exponent or offset during eyes-closed. Lamotrigine reduced corrected theta power in both conditions and alpha power during eyes-open, whereas levetiracetam increased beta power in both conditions and reduced gamma power during eyes-closed.

**Conclusions:** Both lamotrigine and levetiracetam altered aperiodic and periodic EEG activity, with aperiodic effects observed only during eyes-open.

**Significance:** Aperiodic EEG measures may provide a non-invasive approach for characterising ASM-related neurophysiological effects and may help elucidate how distinct pharmacological mechanisms influence large-scale cortical activity.

## 1. Introduction

Electroencephalography (EEG) is a widely used neuroimaging method which captures the summation of the brain’s electrical activity using electrodes placed on the scalp (Biasiucci et al., 2019). EEG signals are comprised of a mix of both periodic (i.e., oscillatory) and aperiodic (i.e., non-oscillatory) neural activity (Donoghue and others, 2020). Early EEG studies highlight the behavioural significance of periodic activity, for example the clear change in the power of oscillations between 8-12 Hz (alpha-band activity) during eyes open and eyes closed resting state EEG recordings (Berger, 1929). Subsequent research defined delta (0.5 – 4 Hz), theta (4-8 Hz), alpha (8-12 Hz), beta (12 – 30 Hz) and gamma (30 – 100 Hz) oscillatory bands, each with distinguishable behavioural and cognitive correlates(Jasper and Andrews, 1938). In addition to periodic oscillatory activity, EEG signals also contain aperiodic non-oscillatory activity, which is represented by the broadband 1/f^β^ background shape of the power spectra (Donoghue and others, 2020; He, 2014). Recent studies have shown that features of the aperiodic 1/f^β^ background, such as the slope steepness (measured by the β exponent) and background offset, can change with different brain states (Lendner et al., 2020), during tasks (Voytek et al., 2015), across the lifespan (Merkin and others, 2023) and in brain disorders (Peterson et al., 2023).

These findings have triggered a renewed interest in understanding the neural origins of aperiodic activity in EEG signals. While the role of synchronous firing of large populations of neurons is well established in generating EEG oscillations, the physiological underpinnings of aperiodic EEG activity are less well understood. Biophysical models of neural activity have posited that the steepness of the spectral slope results from the balance of excitatory and inhibitory postsynaptic potentials mediated by glutamatergic AMPA receptors and GABA_A_ receptors respectively (Gao et al., 2017). In one model, increasing the ratio of excitatory to inhibitory input decreased the spectral exponent (i.e., flattens the slope), whereas increasing inhibitory input increases the exponent (i.e., steepens the slope) (Gao et al., 2017). However, in another model, altering other postsynaptic features such as the decay time of inhibitory postsynaptic current and the leakage current reversal potential also impacted spectral exponents, sometimes in non-linear ways (Brake et al., 2024). In contrast, a simplified single neuron model showed that changes in firing rate resulted in broadband shifts in the offset of the power spectra without impacting the steepness of the slope, particularly over higher frequencies (Miller et al., 2009). Together, predictions from these models suggest that various different neural mechanisms including changes in the ratio of excitation and inhibition could contribute to features of aperiodic EEG activity.

Another method for investigating the neurophysiology of aperiodic EEG activity is to measure changes in EEG power spectra following administration of a drug with known molecular mechanisms of action. In line with predictions from biophysical modelling, drugs that increase inhibition, including GABA_A_ receptor agonists diazepam (Salvatore and others, 2023), pentobarbital (Salvatore and others, 2023), allopregnanolone (Salvatore and others, 2023), propofol (Colombo et al., 2019), and the GABA reuptake inhibitor tiagabine (Muthukumaraswamy and Liley, 2018), increase the spectral exponent (steepen the slope). Furthermore, drugs that decrease excitation, such as perampanel (Muthukumaraswamy and Liley, 2018), a glutamatergic AMPA receptor antagonist, and xenon (Colombo et al., 2019), a glutamatergic NMDA receptor antagonist, also increase the spectral exponent (steepen the slope). Ketamine, another NMDA receptor antagonist which paradoxically increases excitability, possibly via reduced excitatory input to inhibitory interneurons, either has no effect (Salvatore and others, 2023) or decreases the spectral exponent (flattens the slope) (Waschke and others, 2021). However, not all drugs directly support the relationship between excitation/inhibition ratio and spectral slope. For instance, picrotoxin, a GABA_A_ receptor antagonist which decreases inhibition, increases the spectral exponent (steepens the slope) (Salvatore et al., 2024). Furthermore, little is known about how drugs that target other mechanisms which alter neural excitability, such as ion channel blockers or drugs that limit presynaptic neurotransmitter release, impact aperiodic features of EEG activity.

The aim of this study was to investigate how anti-seizure medications (ASMs) which typically reduce neural excitability influence aperiodic EEG activity. We extracted the aperiodic component from resting-state EEG (eyes open and closed) recorded before and after administration of acute doses of lamotrigine (LTG) and levetiracetam (LEV). LTG is a voltage-gated sodium channel blocker which reduces theta oscillations (4-7 Hz), whilst LEV primarily targets synaptic vesicle protein 2A (SV2A) to modulate excitatory neurotransmission which increases beta and gamma oscillations. Beyond these primary mechanisms, LTG also binds nicotinic acetylcholine and 5-HT3 receptors (Vallés et al., 2008), and LEV has been linked to effects on AMPA receptors, monoaminergic and adenosinergic systems, GABAergic transmission, calcium homeostasis, and intracellular pH regulation (Contreras-García et al., 2022). We quantified changes in both the spectral exponent (i.e., slope) and offset of the background of the EEG spectra after accounting for oscillatory peaks using established methods (*specparam*; formerly known as FOOOF) (Donoghue and others, 2020). Given that both drugs are thought to reduce neural excitability, two non-exclusive biologically plausible scenarios were hypothesised: 1) reduced excitability shifts the E/I balance towards inhibition, steepening the slope (i.e., increased exponent); and/or 2) reduced excitability decreases firing rates reducing the broadband offset. We also calculated the spectra after subtracting the aperiodic background to assess whether previously reported oscillatory changes occurred in addition to aperiodic changes following drug intake.

## 2. Methods

### 2.1 Participants

This study utilized data from a previous experiment (Biondi et al., 2022; Premoli et al., 2017). The original study recruited fifteen healthy male volunteers (mean age 25.2 ± 4.62), all of whom provided written informed consent before participating in any experimental procedures. Following transfer and review of the original EEG files for reanalysis, complete data were available for 13 participants, who were included in the current study. All 13 participants had usable data for each drug condition (LTG, LEV, and PBO), eye state (eyes-open and eyes-closed), and pre-and post-administration recording. All participants were confirmed to be right-handed according to the Edinburgh Handedness Inventory (laterality score ≥ 75%). Exclusion criteria included the intake of CNS-active medications, recent use of any drugs (including nicotine and alcohol), and the presence of neurological or psychiatric disorders. Additionally, participants were excluded if they had any contraindications to the medications used in the study (LEV/LTG). The study protocol was approved by the King’s College London Research Ethics Committee (CREC) and conducted in accordance with relevant guidelines and regulations.

### 2.2 EEG Data Acquisition

Participants were seated and instructed to remain silent and still, refraining from any cognitive or mental tasks. EEG data were acquired using a TMS-compatible EEG system (BrainAmp MRplus; Brain Products). The EEG signal was recorded with a 64-electrode EasyCap (EasyCap 64Ch; Brain Products). Electrodes were arranged according to the International 10–20 system, with channel AFz as ground and FCz as reference. The signals were hardware-filtered between DC and 1000 Hz and digitized at a sampling rate of 5 kHz. Conductive gel was applied to each electrode using a blunt-needle syringe to reduce impedance below 10 kΩ, which was maintained throughout the recording sessions.

### 2.3 Experimental Design and Procedure

A single oral dose of either LTG (300 mg), a voltage-sensitive sodium channel blocker, LEV (3000 mg), a specific binder of synaptic vesicle protein 2A, or PBO was administered to participants on separate occasions, each spaced one week apart. This design ensured a sufficient washout period between each condition to prevent any carryover effects. Participants were asked to keep their eyes open and closed in separate conditions to assess different states of brain activity. Resting EEG recordings for both eyes-open and eyes-closed conditions were collected for 3 minutes each and performed at baseline (pre-drug) and 2 hours after drug intake. Concurrent single pulse transcranial magnetic stimulation and EEG conditions were also collected pre and post drug intake separately from the resting EEG recordings but were not analysed in this study and are published elsewhere(Biondi et al., 2022; Premoli et al., 2017). Participants were monitored closely throughout the study to ensure adherence to the protocol and to record any adverse effects. Drug administration and the sequence of eyes-open and eyes-closed conditions varied across sessions to minimize potential biases.

### 2.4 EEG Pre-processing

The raw EEG data files from eyes-open and eyes-closed recordings, both pre-and post-drug administration, were transferred from King’s College London. These files were imported into MATLAB 2022b to undergo a customized pre-processing pipeline using EEGLAB (Delorme and Makeig, 2004). A spatial map of the channel locations was uploaded to the recordings. The data were then down-sampled to 1000 Hz, bandpass filtered between 0.1 and 100 Hz, and notch filtered between 48-52 Hz. Subsequently, the data were segmented into 2-second epochs and pre-and post-drug administration data were concatenated into a single file. Each epoch was visually inspected, and those contaminated with excessive eyeblink or muscle artifacts were removed. Additionally, any disconnected channels (i.e., with flat lines) were identified during this inspection and were excluded. To further refine the data, independent component analysis (ICA) was performed using FastICA. Components associated with residual eye movement and ongoing muscle artifacts were identified and removed following the default criteria of the TESA component selection function (Rogasch et al., 2017). Although some of TESA’s component selection criteria are specific for event-related transcranial magnetic stimulation data (e.g., TMS-evoked muscle activity), the eye blink/movement criteria used here are equally applicable to resting-state data, and the criteria for detecting ongoing muscle activity were developed on resting-state data. Data were then separated back into pre-and post-drug administration time points, missing channels were interpolated, and the data were re-referenced to the average of all electrodes. A summary of epochs retained, independent components removed and channels removed for each condition is presented in supplementary table S1.

### 2.5 Spectral Analysis

Power spectra were calculated using Welch’s method via the PWELCH function in MATLAB. Each channel’s signal was divided into non-overlapping 2-second segments, with the power spectral density (PSD) computed using a Hamming window of the same length. Periodograms were calculated for each segment and averaged to produce the final PSD estimate with a frequency resolution of 0.48 Hz. The computed PSDs and their corresponding frequency values were then saved as JSON files for further analysis.

### 2.6 Spectral parameterisation

The *specparam* analysis (formerly FOOOF) (Donoghue et al., 2020), was conducted in Python and executed in Spyder3 (Anaconda3). The algorithm first estimates the aperiodic component, using the power at the first frequency and the slope between the first and last points of the spectrum to capture the general 1/f power decline. A multi-Gaussian fit then removes oscillatory peaks, isolating the aperiodic component and enhancing the accuracy of the initial fit. The aperiodic exponent and offset were extracted across the frequency range (1-45 Hz) with peak width limits set between 1 and 8, in fixed mode (without a knee). We used the fixed mode parameterisation because the resting-state EEG spectra over the analysed frequency range does not consistently demonstrate a clear bend or knee, as is often evident in invasive recordings. Estimating the knee parameter over a relatively narrow frequency range can reduce the parameter stability and increase the risk of overfitting. The maximum number of peaks, minimum peak height, and peak threshold were set to the default values of unlimited, 0, and 2 SD, respectively. Periodic oscillatory activity was assessed by subtracting the aperiodic component from the original spectra, and then averaging across five frequency bands: delta (2-4 Hz), theta (4-7 Hz), alpha (8-12 Hz), beta (13-30 Hz) and gamma (30-45 Hz). The goodness-of-fit metric, R², was also extracted to assess the explained variance of the model and thus allowing for the assessment of model performance.

### 2.7 Statistical Analysis

To assess changes in aperiodic parameters exponent and offset, periodic frequency bands and goodness of fit, cluster-based permutation tests were conducted using the FieldTrip toolbox in MATLAB (Oostenveld et al., 2011). A within-subjects design was employed, with participant specified as the unit of dependency and condition as the within-subject factor. Statistical comparisons were performed using dependent-samples *t*-tests. Neighbouring channels were defined using a 50-mm Euclidean distance criterion, with clusters required to contain at least two neighbouring channels. An initial cluster-forming threshold of *p* < 0.05 was applied. Cluster-level statistics were calculated using the maximum-sum method, with significance assessed using a Monte Carlo permutation procedure with 5000 randomizations. All tests were two-tailed to detect both positive and negative differences between conditions. Primary outcomes were the change in exponent and offset values (post – pre) following drug conditions (LEV and LTG) compared to placebo during the eyes open and closed conditions. Secondary outcomes measures included change in aperiodic parameters compared to baseline within conditions (e.g., post vs pre), change in corrected periodic oscillatory power for each frequency band following drug conditions (post – pre) compared to placebo, and change in goodness of fit (post vs pre) for each condition.

## 3. Results

### 3.1 The Effect of ASMs on Aperiodic EEG Activity with Eyes Open

#### Offset

First, we examined the effect of ASMs on aperiodic offset within the eyes-open condition (figure 1). LEV did not alter offset values relative to either placebo (PBO) or baseline (no significant clusters for either comparison). In contrast, LTG produced a significant decrease in offset values which was largest over left frontal, central and right parietal electrodes (vs PBO: cluster ∑(t) = −76.7, p = 0.018, d_z_ = −0.86; vs baseline: cluster ∑(t) = −75.1, p = 0.022, dz = −1.03; figure 2).

**Figure 1:**
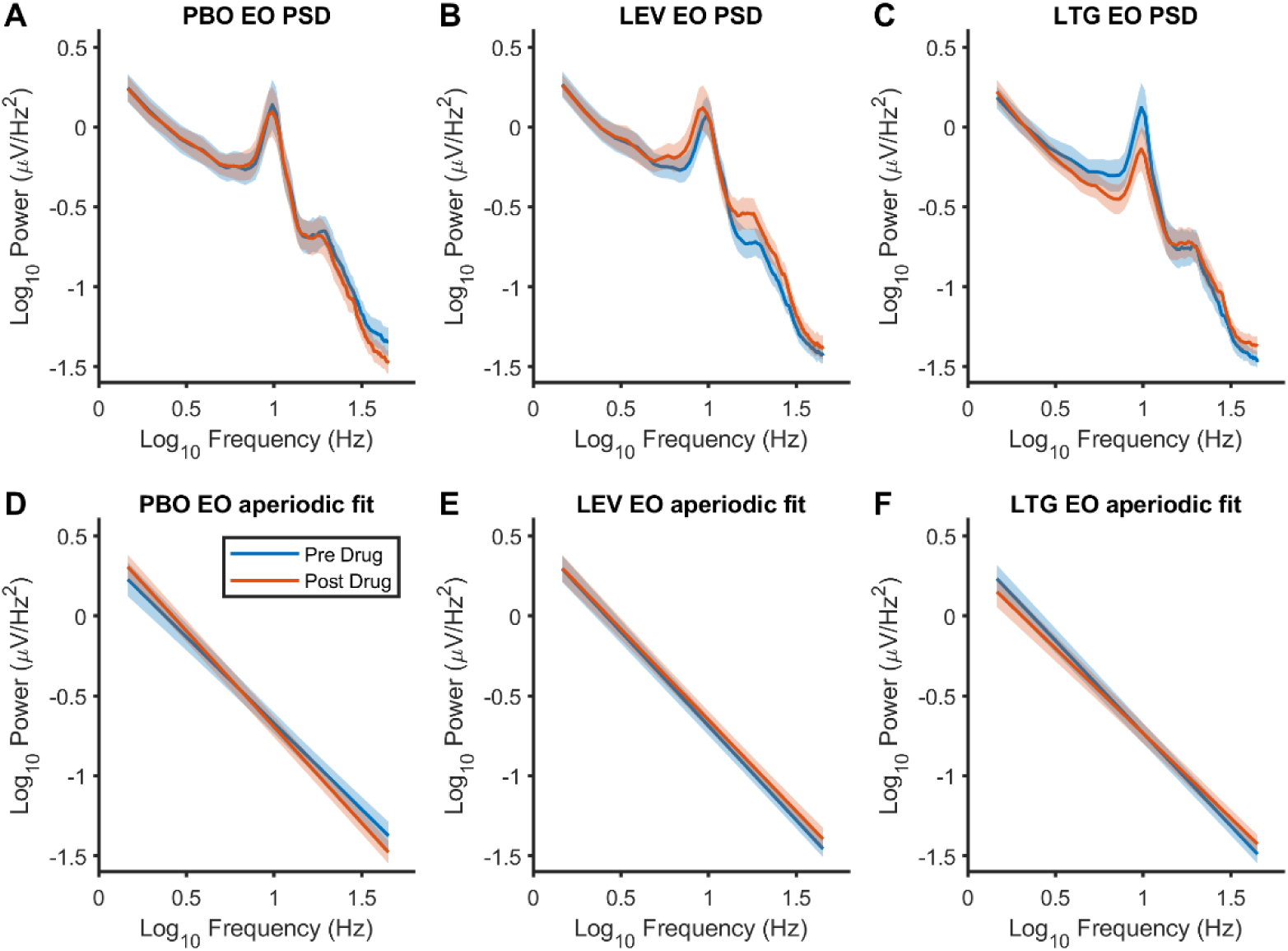
Drug related changes in EEG power spectra and aperiodic fit (eyes open). The top row illustrates the uncorrected power spectral density (PSD) during eyes-open EEG in log-log space for pre- and post-drug conditions. The bottom row shows the corresponding aperiodic model fits. PSD (A-C) and aperiodic model fits (D-F) are averaged across 63 channels. Solid line represents the group mean and shaded bars the standard error of the mean.

**Figure 2:**
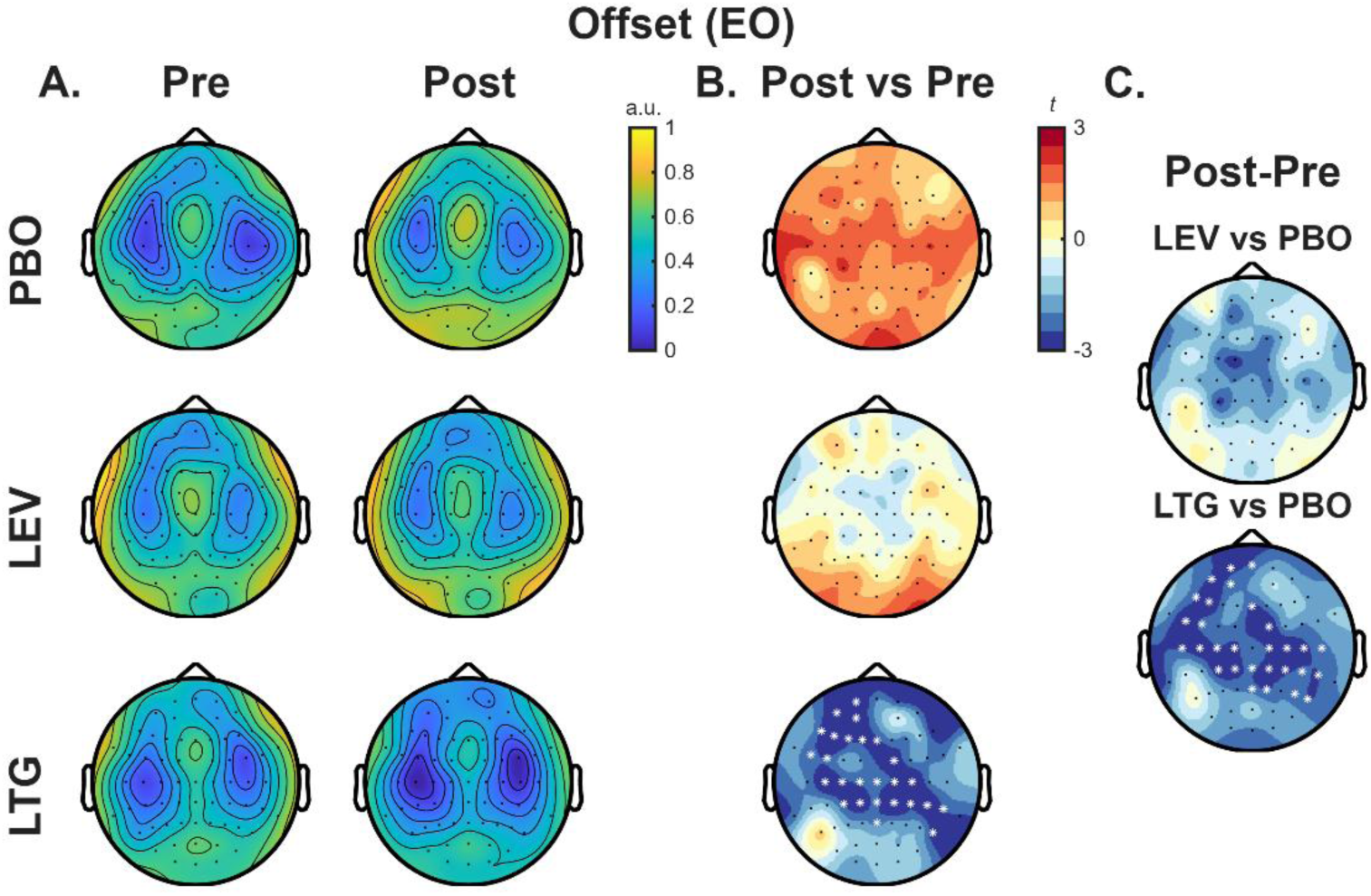
Drug-related changes in aperiodic offset (eyes open). Offset values averaged across participants during the three-minute eyes-open resting state EEG recording. Pre- and post-drug values are shown in (A) and differences between post- and pre-drug offsets presented as t-statistics in (B). The right columns illustrate comparisons of change in offset (post-pre) between drugs and PBO (C). * shows channels contributing to significant clusters (p<0.05).

#### Exponent

Next, we assessed the impact of different ASMs on the aperiodic exponent. LEV did not produce significant pre-to post-drug changes in exponent values (no significant clusters; figure 3). Following LTG administration, two negative clusters were observed over central electrodes (figure 3B) indicating a reduction in exponent values; however neither cluster reached statistical significance (cluster 1: ∑(t) = −10.4, p = 0.094; cluster 2: ∑(t) = −9.8, p = 0.097). In contrast, when comparing the pre-post changes between drug conditions and PBO, both LEV (cluster ∑(t) = −46.0, p = 0.028, d_z_ = −0.82) and LTG (cluster ∑(t) = −41.4, p = 0.045, d_z_ = −0.74) produced significantly greater reductions in exponent than PBO. Figure 4A-B shows individual-subject change plots for both offset and exponent including group means and confidence intervals. Changes in offset and exponent were positively correlated following both LEV and LTG intake (figure 4C-D), suggesting a consistent relationship between changes in these two aperiodic spectral parameters across individuals. Together, these findings indicate that LTG reduced both the aperiodic offset and exponent relative to PBO, whereas LEV selectively reduced the exponent while leaving the offset unchanged during eyes-open resting-state EEG.

**Figure 3:**
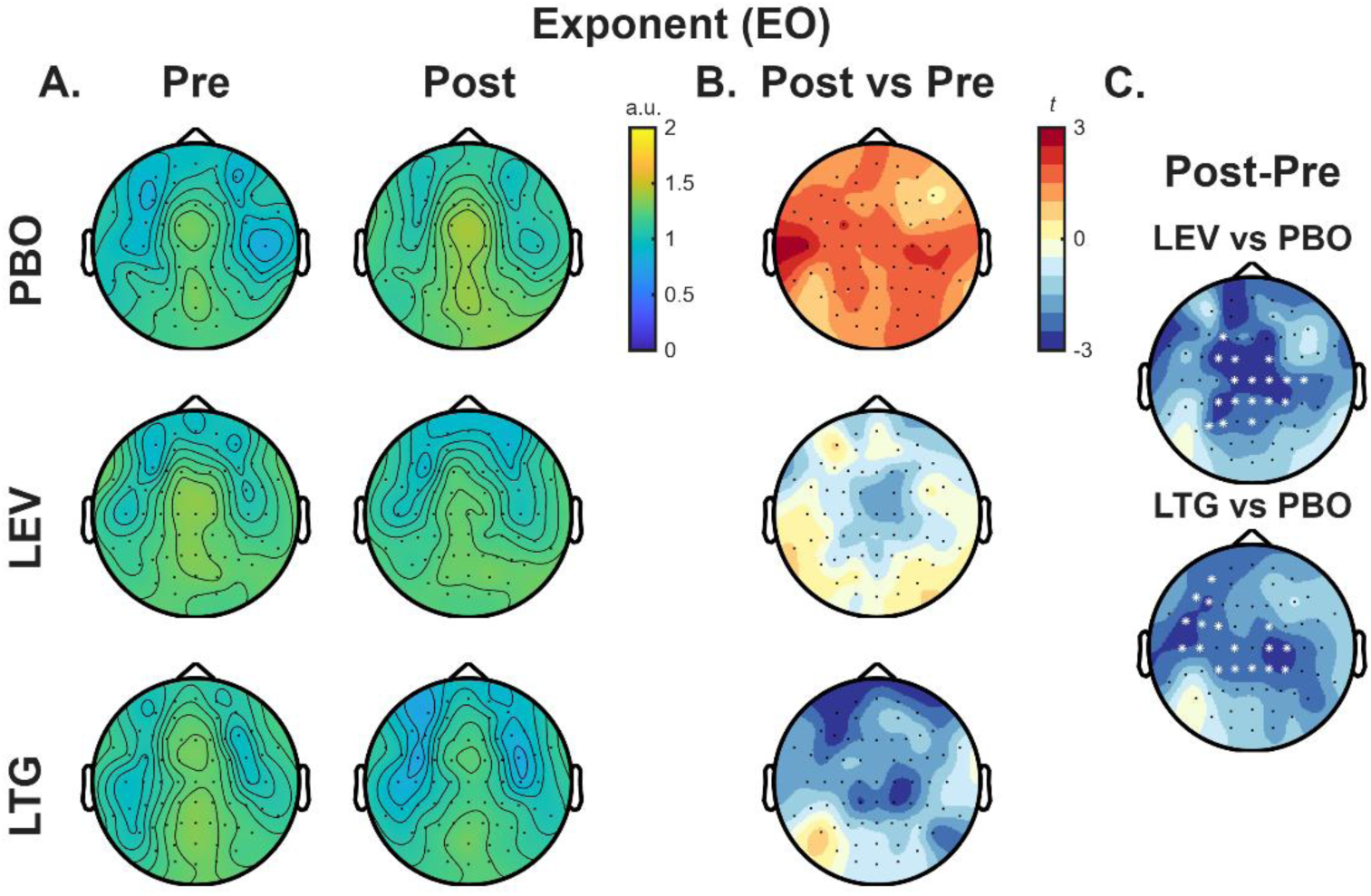
Drug-related changes in aperiodic exponent (eyes open). Aperiodic exponent values during the three-minute eyes-open resting-state EEG recording, averaged across participants. Pre- and post-drug values are shown in (A) and differences between post- and pre-drug values presented as t-statistics in (B). The right columns illustrate comparisons of change in exponent (post-pre) between drugs and PBO (C). * shows channels contributing to significant clusters (p<0.05).

**Figure 4:**
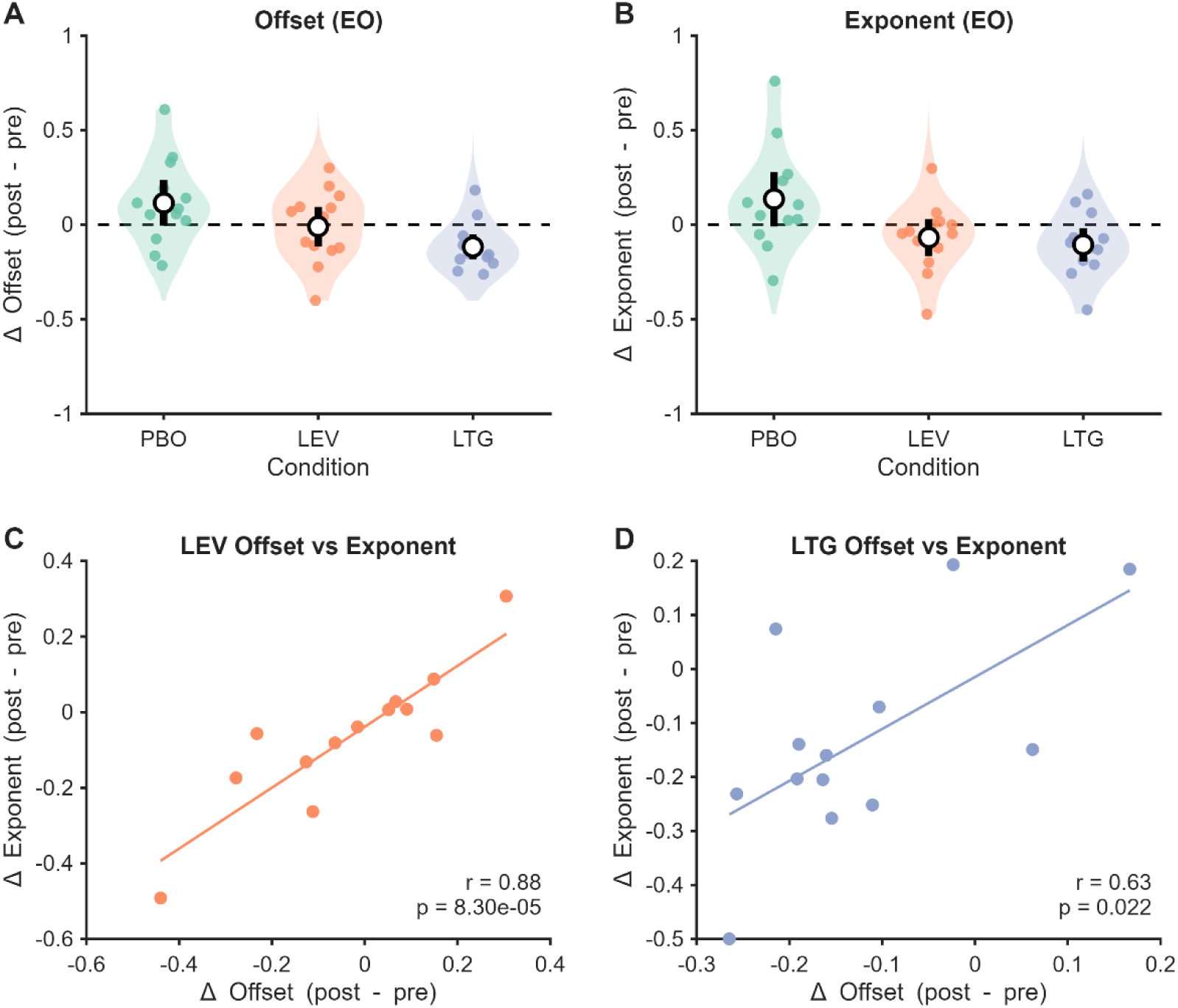
Drug related changes in aperiodic features within individuals. A-B) Coloured data points show the change (post – pre) in offset (A) and exponent (B) for each individual participant following each of the conditions. The white circle shows the group mean and the vertical black bars indicate the 95% confidence intervals. .Horizontal dashed line indicates 0 (i.e., no change). C-D) Coloured data points show the relationship between the change (post – pre) in offset and exponent following LEV (C) and LTG (D) intake. Coloured lines indicate the line of best fit. Correlation coefficients (r; Pearson’s correlation) and p-values are shown in the bottom right corner of each plot

### 3.2 The Effect of ASMs on Aperiodic EEG Activity with Eyes Closed

#### Offset

Figure 5 shows the power spectra before and after drug intake in the eyes closed condition. Neither LEV nor LTG produced significant changes in aperiodic offset value relative to PBO or baseline (no significant clusters; figure 6).

**Figure 5:**
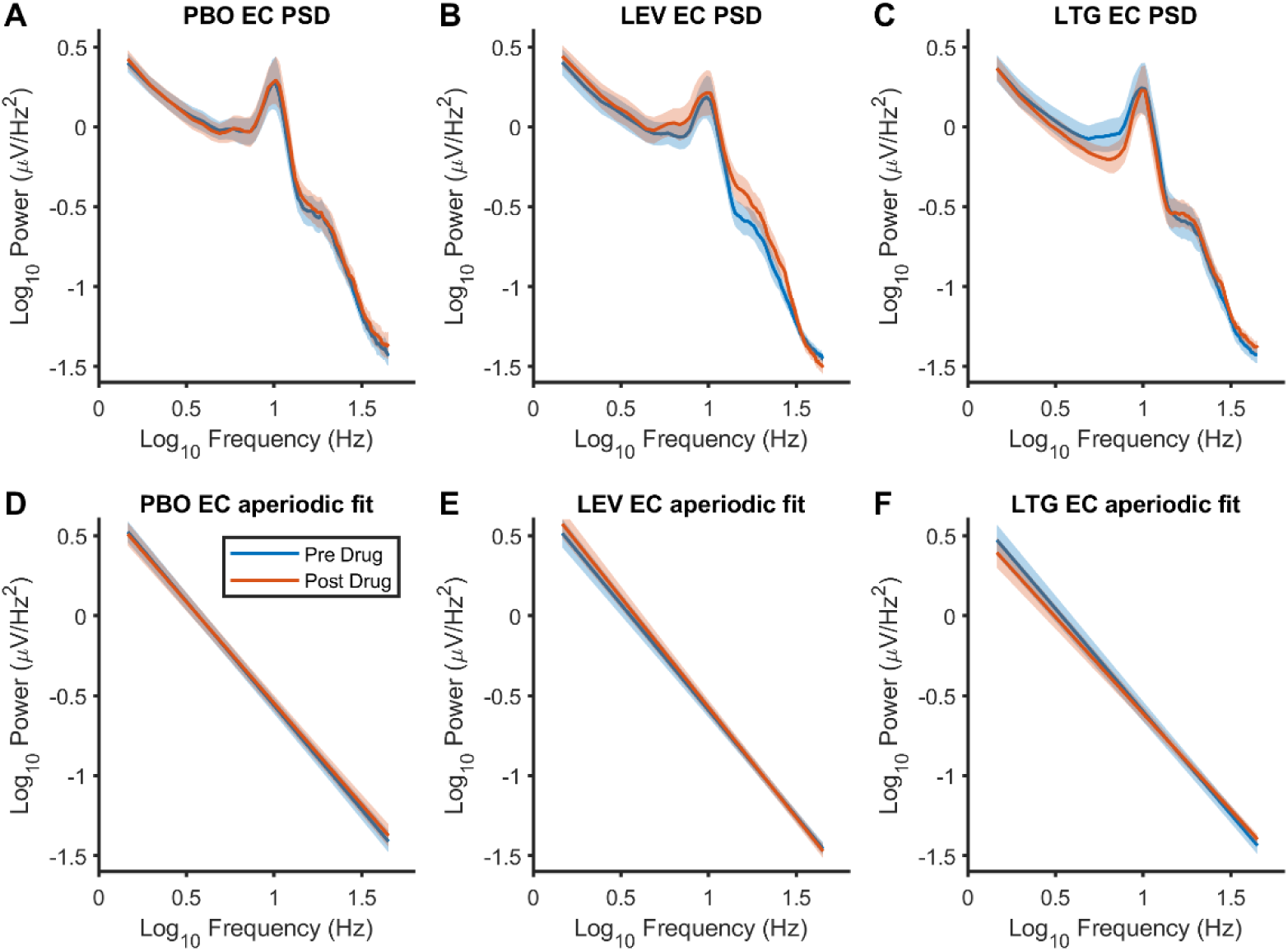
Drug related changes in EEG power spectra and aperiodic fit (eyes closed). The top row illustrates the uncorrected power spectra for pre- and post-drug conditions during the eyes-closed EEG recording, shown in log-log space. The bottom row displays the corresponding aperiodic model fits. Both spectra and aperiodic model fits have been averaged across all 63 channels for each drug condition. Solid line represents the group mean and shaded bars the standard error of the mean.

**Figure 6:**
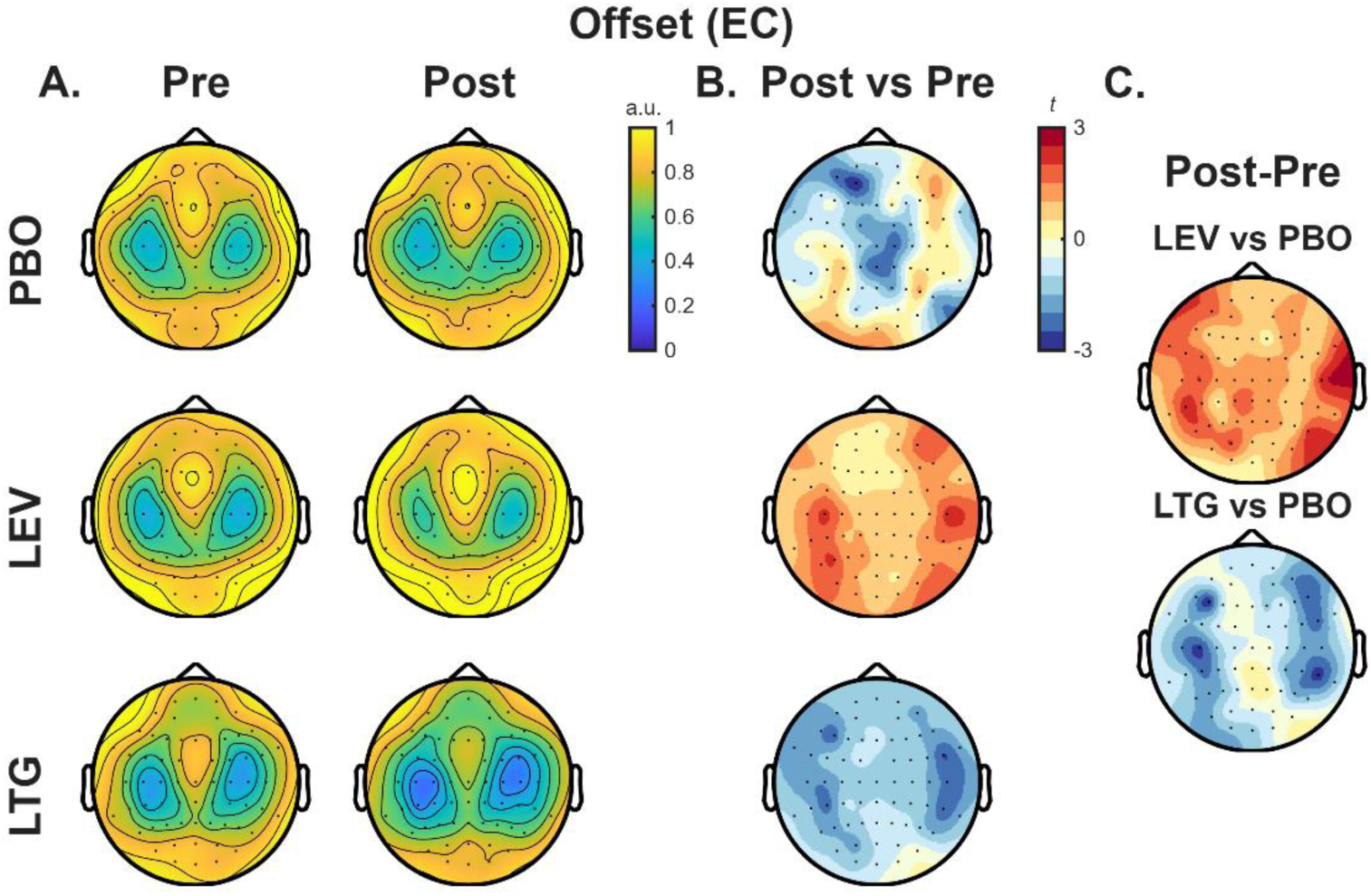
Drug-related changes in aperiodic offset (eyes closed). Offset values averaged across participants during the three-minute eyes-closed resting state EEG recording. Pre- and post-drug values are shown in (A) and differences between post- and pre-drug values presented as t-statistics in (B). The right columns illustrate comparisons of change in offset (post-pre) between drugs and PBO (C).

#### Exponent

Similarly, neither LEV nor LTG altered the aperiodic exponent relative to PBO or baseline (no significant clusters; figure 7). Together, these findings indicate that the reductions in aperiodic offset and exponent observed following ASM administration were specific to the eyes-open condition and were not present during eyes-closed resting-state EEG.

**Figure 7:**
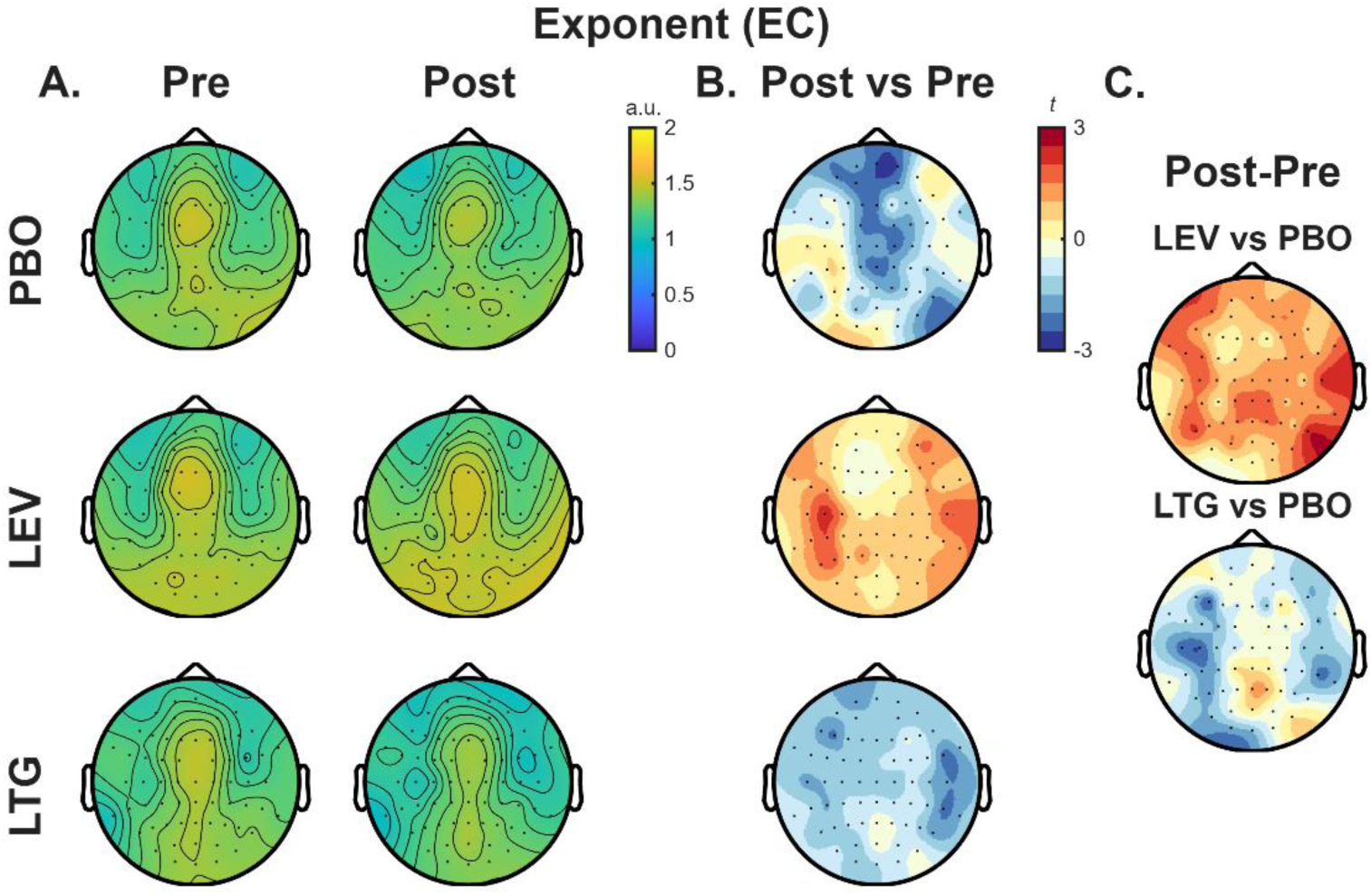
Drug-related changes in Aperiodic Exponent (eyes closed). Aperiodic exponent values during the three-minute eyes-closed resting-state EEG recording, averaged across participants. Pre- and post-drug values are shown in (A) and differences between post- and pre-drug values presented as t-statistics in (B). The right columns illustrate comparisons of change in exponent (post-pre) between drugs and PBO (C)

### 3.3 Sensitivity analysis

To assess the sensitivity of our findings to the choice of fitting range, we repeated the analyses using an alternative range of 2-40 Hz. The results were highly consistent with those obtained using the original 1-45 Hz fitting range, indicating that the observed aperiodic effects were not driven by the specific frequency range selected for parameterization (supplementary figures S1-4).

### 3.4 The Effect of ASMs on Periodic EEG Activity

#### Eyes open

We next assessed changes in periodic activity following ASM intake after correcting for aperiodic activity (secondary outcome) in the eyes open condition. Relative to PBO, LEV produced a significant increase in beta oscillatory power (cluster ∑(t) = 55.5, p = 0.020; d_z_ = 0.97). In contrast, LTG produced significant reduction in theta (cluster ∑(t) = −105.8, p = 0.014, d_z_ = −1.04) and alpha oscillatory power (cluster ∑(t) = −191.2, p = 0.004, d_z_ = −1.08) relative to PBO (figure 8). No significant changes were observed in the remaining oscillatory bands for either drug.

**Figure 8:**
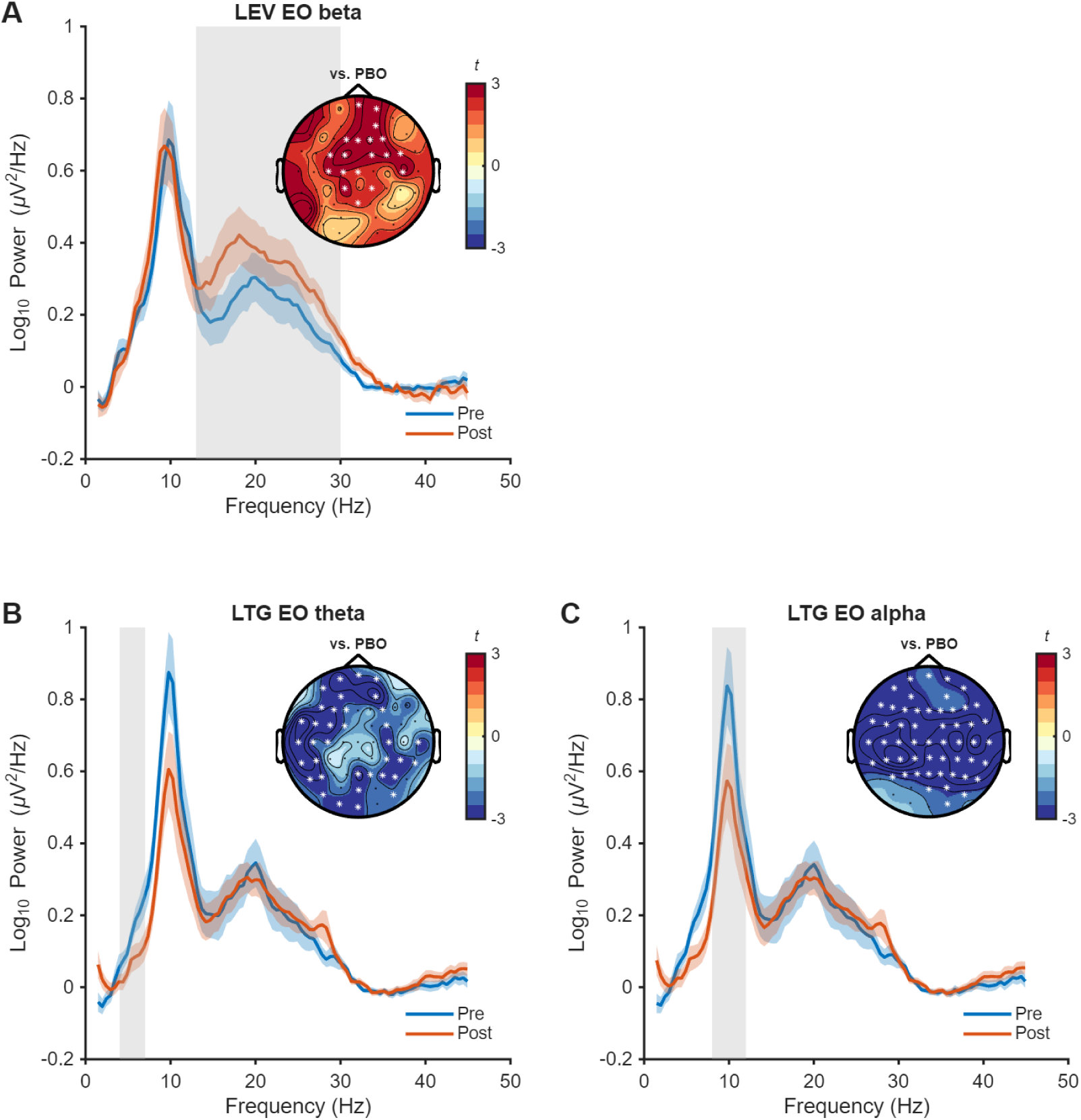
Drug related changes in corrected EEG power spectra (eyes open). Plots illustrate the corrected power spectral density (PSD) after subtraction of the 1/f aperiodic component during eyes-open EEG in semi-log space for pre- and post-drug conditions. Lines show the average of channels contributing to significant clusters and shaded bars show the standard error of the mean. Inset topographies show t-values (drug vs placebo; post-pre for each condition) and significant channels contributing to clusters (*, p<0.05). Shaded grey bars represent the frequency range of significant cluster

#### Eyes closed

In the eyes closed condition, LEV again produced a significant increase in beta oscillatory power (cluster ∑(t) = 155.6, p = 0.004; d_z_ = 1.08) and decrease in gamma power (cluster ∑(t) = −13.3, p = 0.044; d_z_ = −1.36) relative to PBO. LTG produced a significant reduction in theta power (cluster ∑(t) = −114.0, p = 0.005; d_z_ = 1.17; figure 9). No significant differences were identified in the remaining oscillatory bands for either drug. Together, these findings indicate that LEV and LTG modulate periodic oscillatory activity independently of their effects on the aperiodic component.

**Figure 9:**
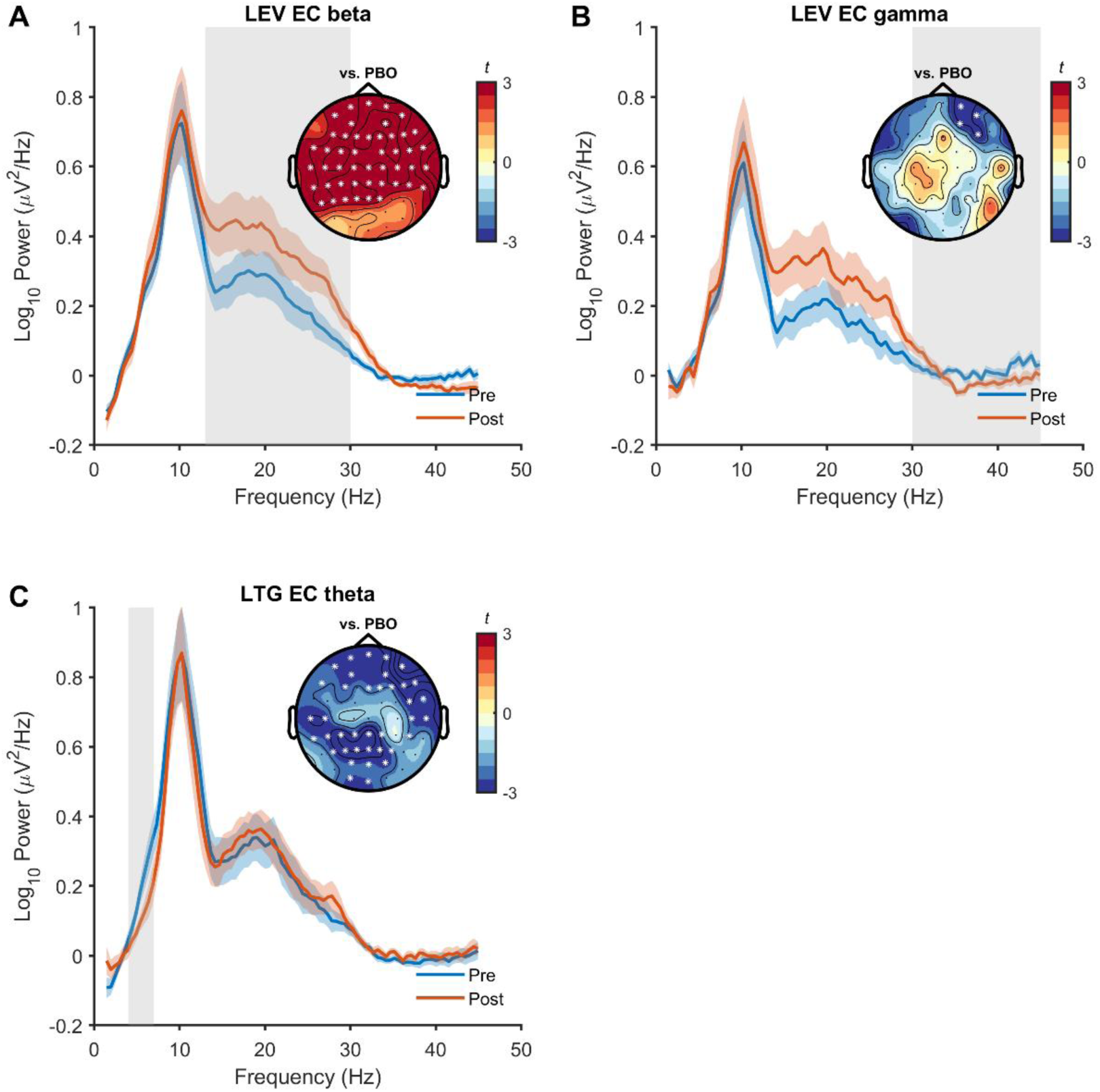
Drug related changes in corrected EEG power spectra (eyes closed). Plots illustrate the corrected power spectral density (PSD) after subtraction of the 1/f aperiodic component during eyes-closed EEG in semi-log space for pre- and post-drug conditions. Lines show the average of channels contributing to significant clusters and shaded bars show the standard error of the mean. Inset topographies show t-values (drug vs placebo; post-pre for each condition) and significant channels contributing to clusters (*, p<0.05). Shaded grey bars represent the frequency range of significant cluster

### 3.5 The effect of drugs on model fit quality

#### Eyes open

Finally, to assess changes in model fit quality over time (control analysis), we extracted the goodness-of-fit parameter (R²) from the *specparam* algorithm. No significant changes were observed following PBO intake (no significant clusters; mean R² ± SD: pre = 0.97 ± 0.05, post = 0.99 ± 0.02, averaged across electrodes) or LTG intake (no significant clusters; pre = 0.98 ± 0.02, post = 0.97 ± 0.06). In contrast, model fit quality significantly declined after LEV intake (max. summed *t* = −8.9, *p* = 0.029; mean R² ± SD: pre = 0.98 ± 0.04, post = 0.98 ± 0.05, Figure 10A), although these changes were numerically small.

**Figure 10.**
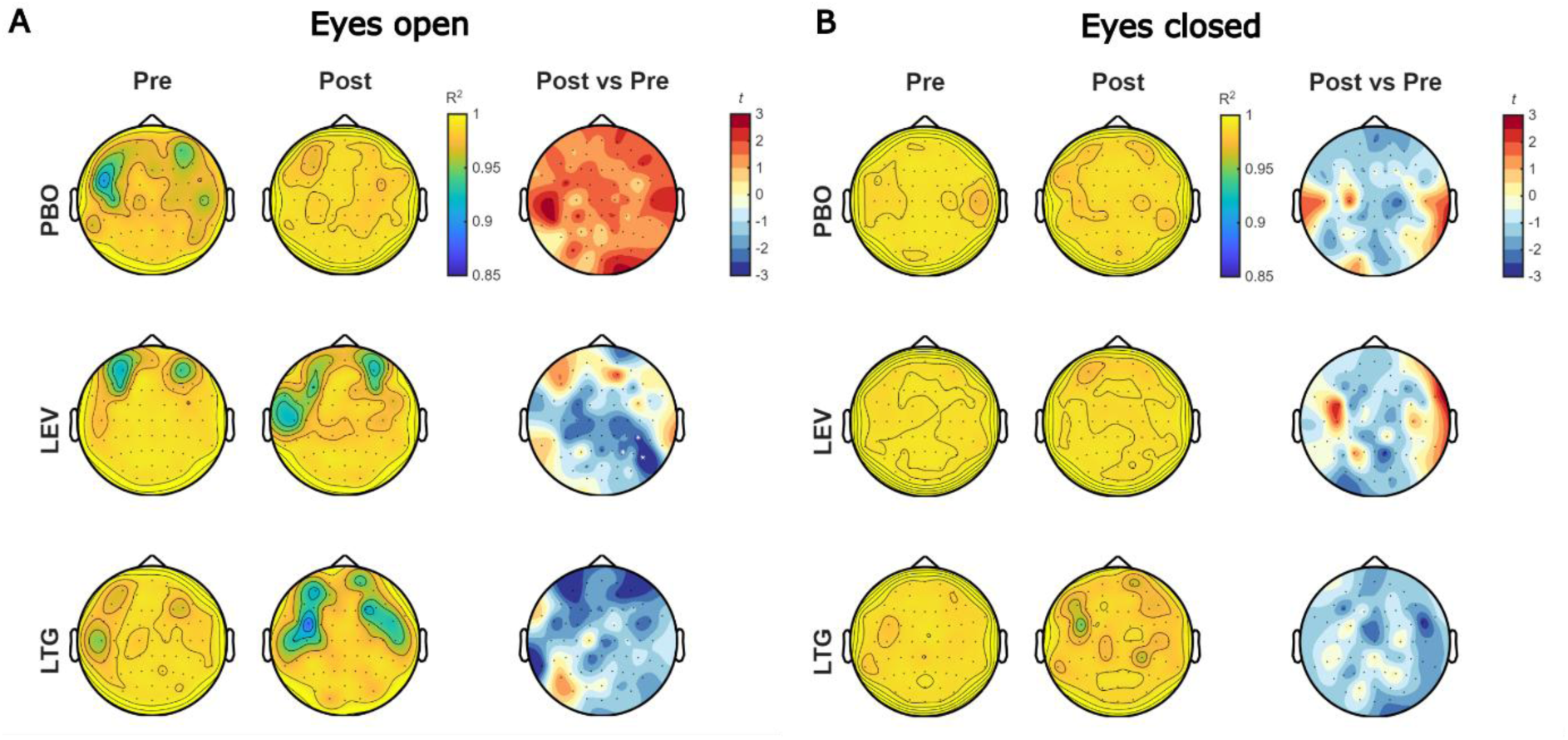
Changes in model fit quality following drug intake. R² values during a three-minute resting-state recording, averaged across participants, for eyes-open (A) and eyes-closed (B) conditions. For each panel, the left and middle column show pre- and post-drug R² values, while the right column displays the difference between post- and pre-drug R² values as t-statistics. * shows channels contributing to significant clusters (p<0.05).

#### Eyes closed

During the eyes closed condition, we could not find evidence for a change in model fit quality following LTG intake (mean R² ± SD; pre = 0.99 ± 0.01, post = 0.98 ± 0.03), LEV intake (mean R² ± SD; pre = 0.99 ± 0.01, post = 0.99 ± 0.01), or PBO intake (mean R² ± SD; pre = 0.99 ± 0.01, post = 0.99 ± 0.01; figure 10B). Together, these findings suggest that changes in model fit quality did not impact changes in aperiodic parameters following drug intake.

## 4. Discussion

The aim of the current study was to extend previous analyses of EEG responses to acute ASM administration by examining changes in aperiodic as well as periodic spectral activity. Using existing data from healthy participants who received LTG, LEV, or placebo, we observed distinct drug-related effects on the aperiodic component of the EEG. During eyes-open recordings, LTG significantly reduced both aperiodic offset and exponent relative to placebo, with the offset reduction largely confined to lower frequencies and the reduced exponent indicating a flattening of the power spectrum. LEV also reduced aperiodic exponent during eyes-open recordings, but did not significantly alter offset. Neither drug significantly altered aperiodic exponent or offset during eyes-closed recordings. In the corrected periodic component, LTG reduced theta power in both conditions and alpha power during eyes-open, whereas LEV increased beta power in both conditions and reduced gamma power during eyes-closed. Overall, these findings extend the previous analyses of the same dataset by demonstrating acute ASM administration can alter both aperiodic and periodic components of EEG activity, with aperiodic changes apparent only during eyes-open recordings and differing between drugs with distinct pharmacological mechanisms.

Lamotrigine (LTG) reduces neuronal excitability by blocking VGSCs and interacting with nicotinic ACh and 5-HT3 receptors (Cunningham and Jones, 2000), effects that likely contribute to its suppression of rapid, sustained firing in response to current injection, as observed in early in vitro studies (Calabresi et al., 1999; Helen et al., 1992; Salvati et al., 1999).Recent research has also demonstrated a notable decrease in the number of action potentials per cell over time following LTG treatment (Kazmierska-Grebowska et al., 2021). In humans, spike rates measured from intracranial electrodes in patients with epilepsy are positively correlated with broadband spectral shifts (e.g., reductions in offset and no change in exponent) (Manning et al., 2009). However, we found a reduction in both the offset and slope in the eyes open condition, suggesting a flattening of the EEG power spectra following LTG. This was accompanied by reductions in the theta oscillatory power in both the eyes open and closed conditions, and alpha oscillatory power decreases specifically in the eyes open condition.

Broadband reductions in spectral power following acute LTG administration align with previous EEG studies assessing longer-term treatment. After a 7-week titration period, Smith *et al*., (2006) reported significant reductions in EEG power between 2-12 Hz, with the most pronounced effects in the 6-12 Hz alpha band. Clemens *et al* (Clemens et al., 2006) expanded upon these findings in a medication-naive epileptic population, reporting reductions in EEG power across the 2-30 Hz range, along with an increase in alpha mean frequency (AMF). Using the same data from the current study, Biondi *et al* (Biondi et al., 2022) reported a more targeted effect of LTG, with a selective reduction in theta band activity during eyes-closed resting-state EEG and no effects in other frequency bands, similar to our current findings taking into account aperiodic activity. Together, these findings suggest that both aperiodic and periodic activity are altered by LTG with a general reduction in power <12 Hz, especially when eyes are open.

Rather than altering VGSC properties (Zona et al., 2001), LEV primarily modulates neurotransmitter release by binding to SV2A (Poulain and Margineanu, 2002). Additional targets such as AMPA, noradrenaline, adenosine, and serotonin receptors, along with mechanisms involving calcium homeostasis, GABAergic transmission, and intracellular pH regulation, have also been reported to contribute to the effects of LEV (Contreras-García et al., 2022). Such molecular actions may underlie findings from single-cell models, which show reduced spike variability and amplitude following LEV administration, likely due to dampened synaptic release (Arato et al., 2021). Reduced spike rates were also observed following LEV administration in another study (Cavitt and Privitera, 2004), which yielded stable offset values, suggesting an alternative underlying mechanism.

In healthy individuals, LEV appears to exert variable influence on EEG power spectra. For example, Mecarelli *et al*., (2004) reported no significant changes in absolute power across any frequency bands during eyes-closed resting-state EEG recordings. However, Salinsky *et al*., (2011) found that in healthy volunteers, LEV administration led to a decrease in lower frequency power, accompanied by increases in alpha and beta bands. Changes have also been observed in patients with epilepsy, although these tend to be limited to higher frequency bands (Park and Kwon, 2009). Using the same data as this study, Biondi *et al*., (2022) observed increases in beta, and gamma power in the eyes-closed condition. We found a reduction in exponent values in the eyes-open condition without a change in offset, suggesting a broadband increase in non-oscillatory higher frequency activity. We also found an increases in beta power after correcting for aperiodic activity in both eyes-open and eyes-closed conditions, suggesting that these alterations reflect genuine oscillatory changes rather than shifts in the aperiodic component. In addition, we found reduced gamma power in the eyes closed condition, however this occurred over lateral frontal channels which are susceptible to muscle artifacts and should be interpreted with caution. Overall, our findings suggest LEV both increases periodic beta oscillatory power and non-oscillatory activity at higher frequencies.

The reasons why LTG-and LEV-induced changes in aperiodic and periodic activity were more pronounced during the eyes-open relative to the eyes-closed condition remain unclear. One possibility is that the findings reflect residual eye-movement-related or myogenic activity that was not removed using ICA. At high doses, LTG has been associated with abnormal slowing of eye movements and drowsiness, and reductions in eye movements can decrease both the aperiodic offset and exponent of resting EEG spectra (Tröndle and Langer, 2026). However, eye-movement-related effects are typically maximal over frontal and occipital regions whereas myogenic activity is maximal over peripheral electrodes overlying or adjacent to cranial and facial musculature (Tröndle and Langer, 2026), which differs from the predominantly centro-parietal distribution observed in the present study. Moreover, abnormal eye movements are generally associated with higher and prolonged LTG dosing, contrasting with the acute single dose administered here. In contrast to the aperiodic findings, the reduction in gamma oscillatory activity observed following LEV intake was predominantly over fronto-lateral channels which may indicate a change in muscle activity. An alternative explanation is that the neurophysiological mechanisms underlying LTG and LEV-induced spectral alterations are more detectable during the eyes-open state, which is characterised by greater sensory input and cortical desynchronisation. Consistent with this possibility, state-dependent changes in EEG power spectra have also been reported for other pharmacological agents, including caffeine(Siepmann and Kirch, 2002), suggesting that stronger effects during eyes-open resting state may not be unique to LTG or LEV. Nonetheless, residual ocular or myogenic contributions cannot be fully excluded and should be considered when interpreting the state-dependent nature of these findings.

Our hypotheses of reduced offset and increased exponent were only partially supported. LTG (VGSC blocker) showed reduced offset and exponent, indicating a reduction in low-frequency aperiodic power and increases in high-frequency aperiodic power, whereas LEV (SV2A modulator) showed a reduced exponent without changes in offset suggesting an increase in high-frequency aperiodic power only. Reductions in low-frequency power are not consistently observed across ASMs that share a primary VGSC mechanism. Carbamazepine (CBZ) acts primarily on VGSCs, with secondary actions at calcium channels and adenosine receptors (Mantegazza et al., 2010), yet has been associated with increased low-frequency power and a reduction in AMF after several weeks of dosing (Marciani et al., 1992). Smith *et al*., (2006) further highlighted these differences by showing that LTG increased AMF, while CBZ did not. Following a single dose of CBZ in healthy individuals, Darmani *et al* (Darmani et al., 2019) also observed an increase in EEG power across multiple bands including theta, alpha and beta frequencies. Following chronic dosing, the increase in power and AMF slowing are thought to result from the neurotoxic impacts of CBZ and may reflect a neurophysiological correlate of impaired cognition (Höller et al., 2018). Together, these findings highlight that drugs with shared molecular mechanisms can result in distinct changes to EEG spectral features, suggesting that multiple mechanisms may contribute to the observed shifts in the EEG spectrum.

Several limitations should be considered when interpreting these findings. First, the sample size was modest and consisted exclusively of healthy male participants, which may have limited sensitivity to detect subtle drug-induced changes and reduces the generalisability of the findings. Second, both drugs were administered as a single acute dose without serum concentration measurements, and therefore the observed effects may differ from those associated with chronic therapeutic administration in epilepsy populations. Third, although extensive preprocessing and ICA-based artifact rejection were applied, residual ocular or myogenic activity cannot be fully excluded, particularly given that LTG and LEV-related effects were observed primarily during the eyes-open condition. Finally, aperiodic parameter estimation is influenced by methodological choices including fitting range, parameterisation approach, and the contribution of oscillatory activity to the spectra. Accordingly, the present findings should be interpreted cautiously until replicated using larger samples and complementary analytical approaches.

## 5. Conclusions

In summary, acute LTG administration was associated with changes in aperiodic and periodic EEG activity in healthy male volunteers. We also observed a reduction in aperiodic exponent relative to placebo following LEV administration, although no significant LEV-related effects were detected for the other aperiodic measures under the conditions examined. Furthermore, changes in aperiodic features were specific to the eyes open condition and not observed with eyes closed for either drug. These findings provide evidence that acute exposure to ASMs can be associated with changes in large-scale neural dynamics reflected in resting-state EEG spectra. However, given the small sample, acute dosing, and absence of clinical or pharmacokinetic measures, these findings should be considered preliminary. Further studies with larger samples, patient populations, chronic dosing, and concurrent measurement of serum drug concentrations are required to establish the robustness, clinical relevance, and potential utility of aperiodic EEG measures for characterising ASM effects.

## 8. Author contributions

**Marissa M. Holden:** Formal analysis, Writing - Original Draft, Visualization. **Isabella Premoli:** Methodology, Investigation, Data Curation, Writing - Review & Editing. **Scott R. Clark:** Writing - Review & Editing, Supervision. **Mark P. Richardson:** Investigation, Writing - Review & Editing. **Mitchell R. Goldsworthy:** Conceptualization, Writing - Review & Editing, Supervision, Project administration. **Nigel C. Rogasch:** Conceptualization, Writing - Review & Editing, Supervision, Project administration

## 10. Declaration of generative AI and AI-assisted technologies in the manuscript preparation process

During the preparation of this work the authors used ChatGPT in order to assist with language editing, improving clarity and structure of the manuscript, and refining the interpretation and discussion of the findings. After using this tool/service, the authors reviewed and edited the content as needed and take full responsibility for the content of the published article.

## 11. Data availability

The dataset analysed during the current study is available from the corresponding author on reasonable request and with permission.

