## Supplementary material for "Investigating the Influence of Anti-Seizure Medications on Aperiodic EEG Activity"

**Supplementary table S1:** Epochs retained, independent components removed, and channels removed during pre-processing.

| | Eyes open | | | | | |
|---|---|---|---|---|---|---|
| | PBO | | LEV | | LTG | |
| | Pre | Post | Pre | Post | Pre | Post |
| Epochs retained | 88.0 (3.3) | 87.6 (3.0) | 87.9 (3.8) | 84.3 (8.5) | 87.7 (3.9) | 87.8 (4.4) |
| ICs removed | 20.0 (13.9) | | 22.8 (12.7) | | 21.2 (11.1) | |
| Channels removed | 0.0 (0.0) | | 0.2 (0.6) | | 0.0 (0.0) | |
| | Eyes closed | | | | | |
| Epochs retained | 88.5 (4.8) | 89.3 (1.6) | 88.9 (2.5) | 87.8 (2.0) | 88.9 (1.3) | 87.7 (3.8) |
| ICs removed | 21.5 (12.4) | | 22.0 (11.2) | | 21.7 (12.0) | |
| Channels removed | 0.0 (0.0) | | 0.2 (0.6) | | 0.0 (0.0) | |

Data = mean (± standard deviation). IC = independent component.

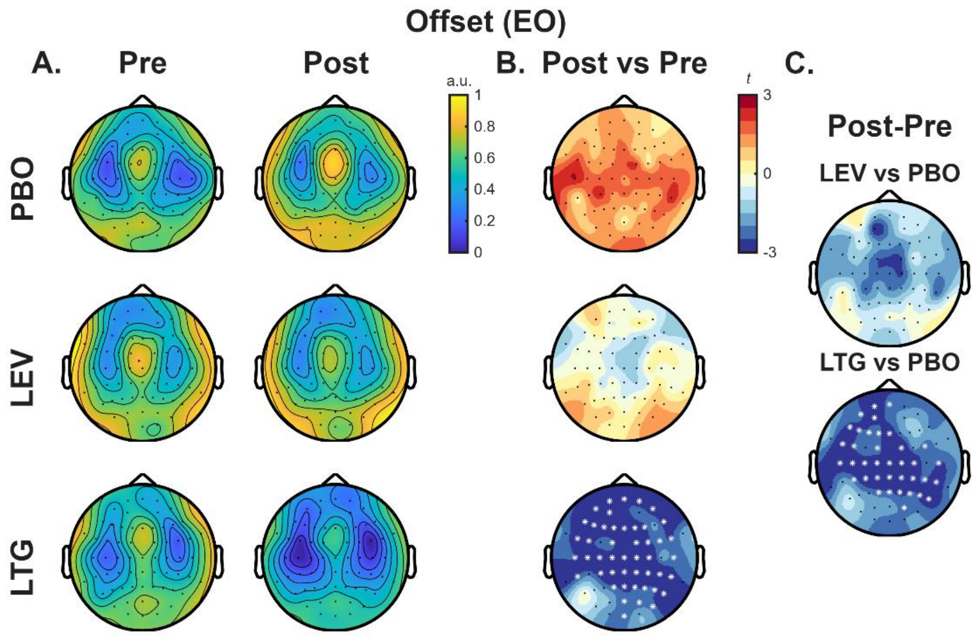

***Figure S1: Drug-related changes in aperiodic offset using 2-40 Hz fitting range (eyes open).*** *Offset values averaged across participants during the three-minute eyes-open resting state EEG recording. Pre- and post-drug values are shown in (A) and differences between post- and pre-drug offsets presented as t-statistics in (B). The right columns illustrate comparisons of change in offset (post-pre) between drugs and PBO (C). * shows channels contributing to significant clusters (p<0.05).*

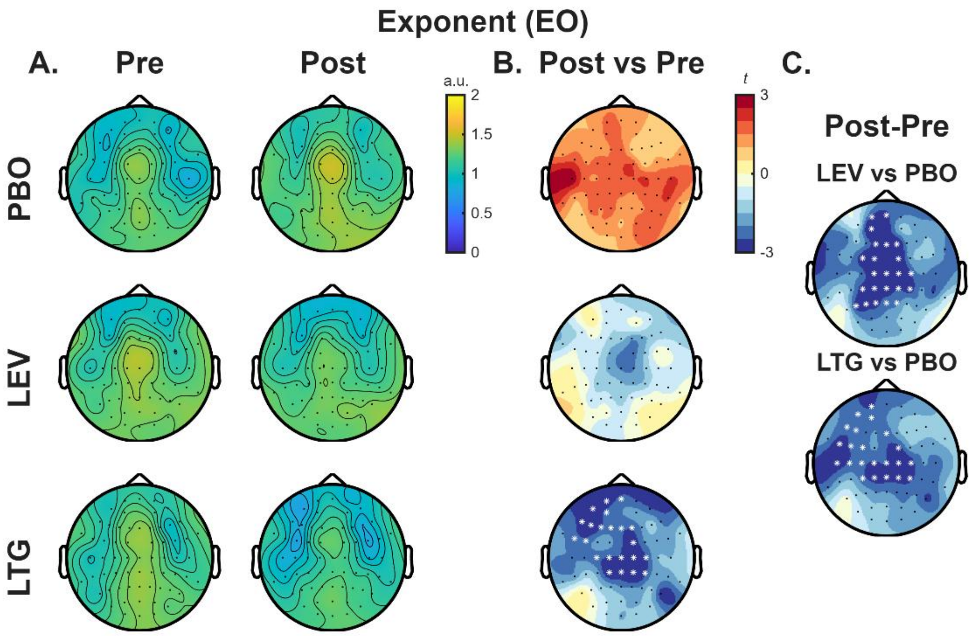

***Figure S2: Drug-related changes in aperiodic exponent using 2-40 Hz fitting range (eyes open).*** *Aperiodic exponent values during the three-minute eyes-open resting-state EEG recording, averaged across participants. Pre- and post-drug values are shown in (A) and differences between post- and pre-drug values presented as t-statistics in (B). The right columns illustrate comparisons of change in exponent (post-pre) between drugs and PBO (C). * shows channels contributing to significant clusters (p<0.05).*

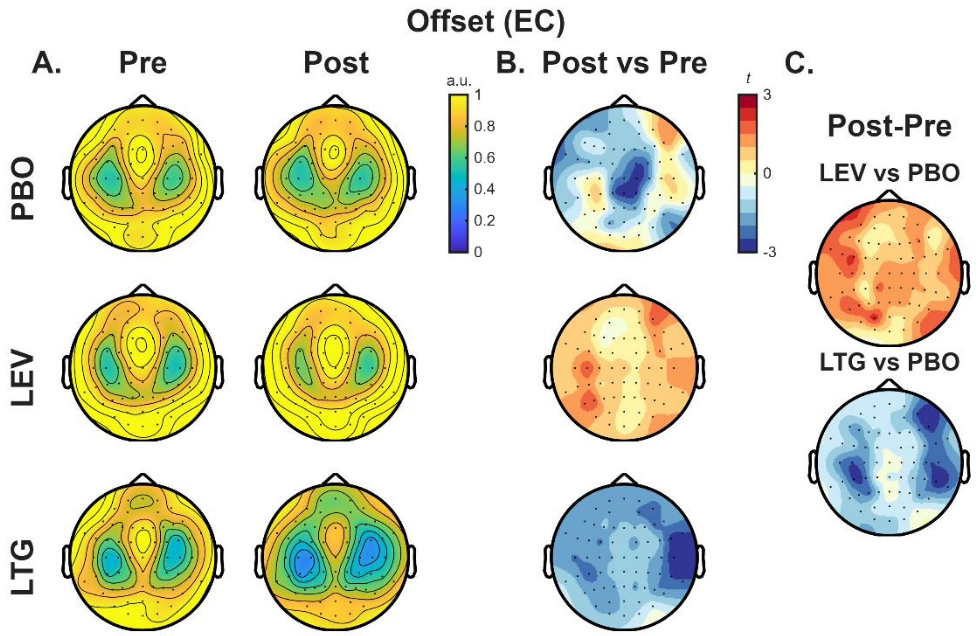

***Figure S3: Drug-related changes in aperiodic offset using 2-40 Hz fitting range (eyes closed).*** *Offset values averaged across participants during the three-minute eyes-closed resting state EEG recording. Pre- and post-drug values are shown in (A) and differences between post- and pre-drug values presented as t-statistics in (B). The right columns illustrate comparisons of change in offset (post-pre) between drugs and PBO (C).*

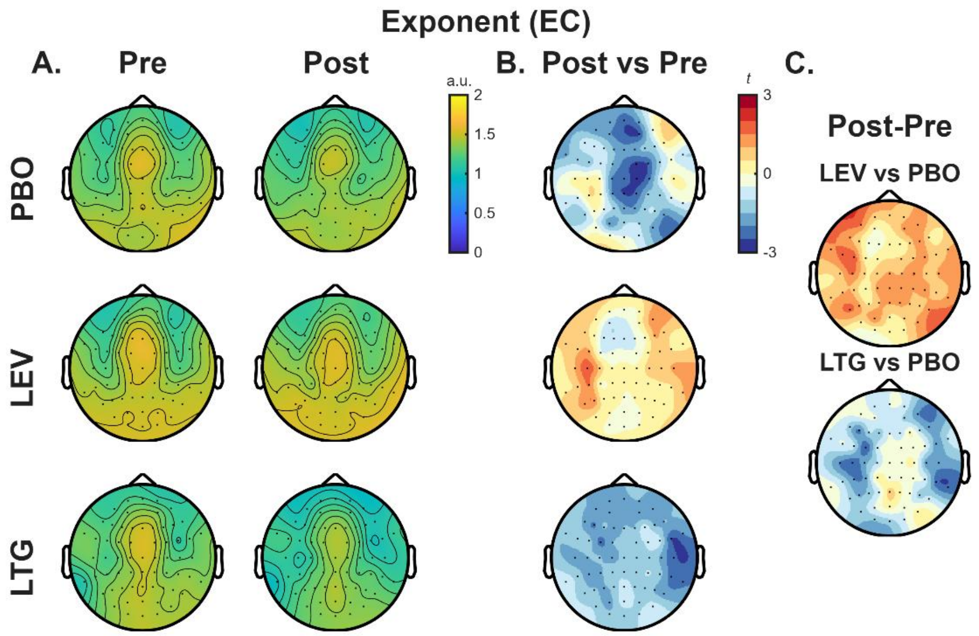

***Figure S4: Drug-related changes in Aperiodic Exponent using 2-40 Hz fitting range (eyes closed).*** *Aperiodic exponent values during the three-minute eyes-closed resting-state EEG recording, averaged across participants. Pre- and post-drug values are shown in (A) and differences between post- and pre-drug values presented as t-statistics in (B). The right columns illustrate comparisons of change in exponent (post-pre) between drugs and PBO (C).*
